# Bittersweet dynamics from flowers to fruits: chemometric molecular networking reveals metabolic changes towards reduced toxicity over ontogeny in above-ground *Solanum dulcamara* chemotypes

**DOI:** 10.64898/2026.07.29.741467

**Authors:** Redouan Adam Anaia, Ilaria Chiocchio, Nicole M. van Dam

**Affiliations:** Groningen Institute for Evolutionary Life Sciences, University of Groningen, Nijenborgh 7, 9747 AG, Groningen, The Netherlands; German Centre for Integrative Biodiversity Research (iDiv) Halle-Jena-Leipzig; Leibniz Institute for Horticultural Sciences (IGZ), Großbeeren, Germany; Department of Pharmacy and Biotechnology, University of Bologna, Via Gobetti 87, 40129 Bologna, Italy; Institute of Biodiversity, Ecology and Evolution (IBEE), Friedrich Schiller University, Jena, Germany

**Keywords:** metabolome mining, steroidal glycosides, phenolamides, hydroxycinnamates, acetylation, acylation, LC-MS, untargeted metabolomics

## Abstract

Poisonous plants frequently deploy toxic metabolites in an organ- and ontogenetic-specific manner, yet the developmental dynamics of these plant specialised metabolites remain poorly understood. In the Solanaceae, steroidal glycosides (SGs), including steroidal glycoalkaloids (SGAs) and steroidal saponin glycosides (SSGs), are major defence metabolites with potent ecological and pharmacological activities. Here, we investigated how their metabolic profiles are reorganised during flower and fruit development in *Solanum dulcamara* using untargeted metabolomics and chemometric molecular networking.

Non-metric multidimensional scaling (NMDS) using classified metabolite features recovered clear organ-specific and chemotype-specific segregation of samples, which was further strengthened when analyses were restricted to metabolomic features annotated as SGs. In flowers, orthogonal projections to latent structures discriminant analysis (OPLS-DA) separated both floral developmental stage and chemotype, demonstrating that developmental stage and inherited SG polymorphism represent independent axes of chemodiversity in flowers. Chemometric molecular networks showed pronounced diversification of hydroxycinnamate metabolism towards anthesis, including accumulation of caffeoylputrescine in flower buds and progressive acylation of spermidine conjugates across flower development. Upon anthesis, glycosylated hydroxycinnamates and flavonoids, and hydroxycinnamoyl-acylated flavonoid glycosides are accumulated. Fruit ripening followed a contrasting trajectory, in which SGAs were associated with unripe fruit pericarp, whereas ripe pericarp accumulated N- and O-acetylated SGA derivatives, oxidative deamination products, and predicted steroid degradation products, indicating stepwise detoxification of toxic steroidal glycosides during seed maturation.

Together, these findings show that the poisonous chemistry in *S. dulcamara* is highly dynamic and developmentally orchestrated. Flowers accumulate structurally complex metabolites toward anthesis, including conjugates of hydroxycinnamates, whereas fruits metabolise toxic SGAs into less toxic SGs during ripening, linking metabolic remodelling to reproduction and seed dispersal.

## Introduction

The genus Solanum is the largest genus in the Solanaceae comprising half of the species in this family. Next to numerous wild species, it also gave rise to several crops, such as tomato (*Solanum lycopersicum*), eggplant (*S. melongena*) and potato (*S. tuberosum*) (Särkinen et al., 2013). Phytochemically, the genus *Solanum* is characterized by the biosynthesis of a group of cholesterol-derived plant specialized metabolites (PSMs), collectively called steroidal glycosides (SGs; Eich, 2008). Within this class of metabolites, the chemistry and biosynthesis of steroidal glycoalkaloids (SGAs) are well studied, due to their toxicity to humans and for providing resistance to pathogens and herbivores to the plant (Cárdenas et al., 2015; Sonawane et al., 2018; Wolters et al., 2023). A series of Glycoalkaloid Metabolism (GAME) genes involved in the biosynthesis of SGAs as well as in that of structurally related steroidal saponin glycosides (SSGs) has been identified in tomato and potato (Boccia et al., 2024; Cárdenas et al., 2015; Itkin et al., 2013; Sonawane et al., 2017, 2020).

Leaves, roots and fruits (berries) of *Solanum dulcamara*, a wild Eurasian nightshade species also known as Bittersweet nightshade, predominantly accumulate SGAs, and to a lesser extent SSGs (Anaia et al., 2025; Calf et al., 2018; Eich, 2008; Köthe & Willuhn, 1983; Sander & Willuhn, 1961; Willuhn, 1967, 1968, 1969; Willuhn & Kun–anake, 1970). SGs from *S. dulcamara* are of ethnobotanical importance, as different plant organs have been traditionally used for their pharmacological activities (Asfaw et al., 2023; Gafforov et al., 2024; Kaunda & Zhang, 2019). Multiple naturally occurring and heritable SGA leaf chemotypes with different levels of resistance to specialist and generalist herbivores have been described (Calf, 2019). In *S. dulcamara* plants, SGAs serve as defence compounds to slugs, which are generalist herbivores (Calf et al., 2018, 2019). Specialist herbivores, in particular flea beetles, fed more on plants containing SGAs, whereas an accession containing mainly SSGs conjugated with uronic acids was avoided (Calf et al., 2019). This indicates that SGAs may serve as cues or feeding stimulants to adapted herbivores like flea beetles.

One specific chemical SG polymorphism in *S. dulcamara* is based on the level-of-unsaturation, or ring-double bond equivalent (RDBE) of the steroidal moiety (Anaia et al., 2025). In tomato, the conversion from a unsaturated to a saturated SGA leaf chemotype is associated with the expression of *GAME25*, encoding a short-chain dehydrogenase/reductase (Sonawane et al., 2018). In *S. dulcamara*, the expression levels of a homologue, *SdGAME25,* were higher in the leaves of vegetative plants with saturated SGs (S-chemotypes) than in those with mainly unsaturated SGAs (U-chemotypes). However, both the levels of saturated and unsaturated SGs and the expression of GAME25 were found to differ among leaves and roots as well as over ontogeny (Anaia et al., 2025). The root metabolomes of different *S. dulcamara* leaf chemotypes were less distinct, though there are differences in SG profiles among embryonic and adventitious roots (Anaia et al., 2025; Chiocchio et al., 2023).

The rather small differences in the steroidal aglycon structure of *S. dulcamara* chemotypes were found to have ecological effects. In a common garden experiment, *S. dulcamara* plants of the U-chemotype suffered more damage by leaf chewing herbivores, mainly *Leptinotarsa decemlineata* than S-chemotypes (Anaia & van Dam, *preprint*). At the same time, flowers of plants with U-chemotypes were visited more often by *Bombus lapidarius* bumble bees than flowers of plants with the S-chemotype (Anaia & van Dam, *preprint*). This suggest that not only herbivores, but also pollinators may perceive chemical differences in flower or pollen of different *S. dulcamara* chemotypes. It is likely that flower and leaf metabolomes are phytochemically and biosynthetically similar to each other, as flowers originate from the same meristems as leaves, only with a different, genetically determined developmental program (Ratcliffe et al., 1998). The differences between S- and U-chemotypes may also be visible in the fruits. The pericarp of solanaceous fruits is formed from the ovary wall, which is maternal tissue. However, the chemical composition of fruits may differ over time. Analyses of tomato fruits showed that during ripening, the fruit metabolome changes (Martínez-Rivas & Fernie, 2024). A series of enzymes and a transporter convert the bitter SGA α-tomatine in green tomatoes to the non-bitter and less toxic esculeoside A in red tomatoes (Akiyama et al., 2021; Kazachkova et al., 2021; Sonawane et al., 2023; S. Wang et al., 2023; Yamanaka et al., 2009). Not only makes this commercial tomatoes palatable for human consumption, but it also makes the fruits of wild Solanaceae, such as *S. dulcamara*, more attractive and palatable to seed dispersers, such as birds (Cipollini & Levey, 1997b; Symon, 1979; Whitehead et al., 2022). Whether the fruit ripening process may make initial differences among leaf and fruit chemotypes less distinct, is as yet unknown.

Using untargeted metabolomics, we tested whether the SG-based leaf metabolic profiles of U- and S-*S. dulcamara* chemotypes are also reflected in their flowers and fruits. Therefore, we sampled flowers and fruit pericarp at different developmental stages to analyse whether their metabolomes may change and become more similar during floral development and fruit ripening, respectively. In addition, we analysed root and leaf metabolomic profiles. Based on the extensive body of tomato research, we hypothesized that flower and pericarp chemotypes align with leaf chemotype, but not with root metabolic profiles. Moreover, we tested the hypothesis that the metabolic profile of pericarp of green berries is more similar to leaf metabolic profiles than that of red berries. Finally, we expected that root metabolic profiles were less distinct among chemotypes that those of leaf, flowers or pericarp. We sampled roots, leaves, and flowers as well as unripe and ripe berries from the F_1_ of two well described chemotypes with different leaf SGA profiles. We used offspring from two accessions with known leaf chemotypes: ZD04 which is a plant of U-chemotype, and TW12 which is a plant with S-chemotype (Calf, 2019; Calf et al., 2018). By creating selfings of either parent and crosses between the two, we generated a F_1_ and two S_1_ populations segregating for SGA chemotypes (Calf, 2019; Willuhn, 1966). After extraction and chemical analyses, we used multivariate analyses combined with computational metabolomics to compare their metabolic profiles. Thereby, we aimed to explore the phytochemical space of semi-polar metabolites in *S. dulcamara* to connect the metabolome to previously observed chemotypes (Vitale et al., 2024).

Our analyses revealed that flower and unripe fruit SG chemotypes are indeed more similar to their respective leaf chemotype than to their root chemical profile. Flowers and unripe fruits have SG chemotypes that resemble those of leaves, hinting towards co-regulation of SG biosynthesis in above-ground organs. However, among the reproductive organs there were additional levels of phytochemical diversification. Flowers were especially enriched in SSGs, while berries predominantly contained N-and O-acetylated SGAs. Additionally, we found that acylation of polyamines and flavonoid glycosides with hydroxycinnamic acids increases with the progression of floral development in *S. dulcamara.* Lastly, during fruit ripening, we found that the toxic *S. dulcamara* SGAs present in green fruits were acetylated and degraded in ripe fruits. Together, our data indicated that there is multi-layered chemodiversity in present *S. dulcamara,* which changes across organ development to optimize biotic interactions.

## MATERIALS AND METHODS

### Plant material

Seeds from three *Solanum dulcamara* crosses (TW12 S_1_, TW12 x ZD04 F1 and ZD04 S_1,_ Table 1) which segregate for SG chemotype (Calf, 2019) were germinated after two weeks of cold stratification at 4 °C. The seedlings were transplanted to trays (QuickPot™ 24R, Ø 7.5 × 7.0 cm; Groß Kreutz, Germany) filled with a 1:1 (v/v) autoclaved soil (Floradur B pot clay medium coarse; Floragard Vetriebs, Germany) and sand (0/2 washed; Rösl Rohstoffe, Germany) mixture and grown for 4 weeks under greenhouse conditions (17–25 °C, RH: ±65%) with light supplemented to 280 μmol·m^−2^·s^−1^ with high-pressure sodium lamps (Anaia et al., 2025). Then, plants were potted in 1 L pots (11 × 11 × 12 cm) using the aforementioned soil:sand mixture supplemented with 4 g·L^−1^ Osmocote Pro 8-9M (ICL Boulby, Cleveland, UK). Plants were watered every other day, and additional liquid fertilizer was supplied whenever needed. In total, 174 plants were grown and samples were taken across plant ontogeny to ensure that plants of contrasting ontogenetic stages were sampled as plant ontogeny might affect SG profiles (Anaia et al., 2025). Harvest was started when the first plants showed transitioning meristems and continued until all samples were taken. At this time, a *Bombus terrestris* hive (BioBest Nederland B.V., De Lier, The Netherlands) was introduced into the greenhouse to facilitate buzz-pollination of *S. dulcamara* flowers. Different organs of a same plant were harvested consecutively, thereby prioritizing above-ground organs. Flower buds (hereafter “pre-flower”), closed and open flowers were sampled in whole (including calyx, anthers, pollen and stamen) by cutting at the base of the flower. The pericarp and placental tissues, together referred to as ‘pulp’ hereafter, of the berries were sampled by cutting the fruits in half with a scalpel and quickly removing the seeds. For above-ground organs, samples were directly flash-frozen in liquid nitrogen upon harvest, ground with mortar and pestle under liquid nitrogen, and were kept at −80 °C until they were freeze-dried. For harvests of taproots, roots were washed on a root washing table using a pressure hose after removal from the pots. Then the roots were gently dried using tissue paper, harvested using scissors and put in 50 mL Falcon tubes, which were flash-frozen in liquid nitrogen. Then, roots were freeze-dried and ground into a fine powder using a ball mill (Mixer Mill MM 400; Retsch GmbH, Haan, Germany). In total, we took the following samples: leaves (n=180), taproot (n=159), pre- (n=104), closed (n=73) and open flowers (n=78), pericarps of unripe (green, n=28) and ripe (red, n=34) berries of *S. dulcamara*.

**Table 1:** *Solanum dulcamara* crosses used for profiling of semi-polar metabolites and expected steroidal glycoside (SG) inheritance pattern. Abbreviations - SGAs: steroidal glycoalkaloids; SSGs: steroidal saponin glycosides; TW: Texel Wet; ZD: Zandvoort Dry; *U*: unsaturated; *S*: saturated;. Parental accessions TW12 and ZD04 predominantly accumulate *S*-SGAs and *U*-SGAs in leaves, respectively (Calf, 2019; Calf et al., 2018).

| Accession | Recipient ♀ | Donor ♂ | Individuals | Expected |  |
| --- | --- | --- | --- | --- | --- |
|  |  |  |  | SG chemotypes | Ratio |
| TW12 S1 | TW12 | TW12 | 38 | S-SGAs : U-SGAs | 3:1 |
| ZD04 S1 | ZD04 | ZD04 | 32 | U-SGAs : U-SSGs | 3:1 |
| TW12 x ZD04 F1 | TW12 | ZD04 | 104 | S-SGAs : U-SGAs | 1:1 |

### Extraction of endogenous semi-polar metabolites

For extractions of semi-polar metabolites, 20.0 mg of freeze-dried, powdered material was weighed into an 2.0 mL Eppendorf tube and extracted using 7:3 [v/v] methanol:water (Anaia et al., 2025; Chiocchio et al., 2023). We used a procedure derived from De Vos et al., 2007 combined with Rogachev & Aharoni, 2012. Briefly, an extraction buffer solution was prepared by mixing acetate buffer (2.3 mL acetic acid and 3.41 g ammonium acetate added to 1 L millipore water at pH 4.8) with MS-grade methanol in a 1:3 ratio. The extraction buffer was kept at approximately 4 °C, and 1 mL extraction buffer was added to 2 mL Eppendorf tubes containing two metal beads (3 mm Ø) and 20 mg of ground freeze-dried leaf tissue. Eppendorf tubes were shaken for 10 seconds at 30 Hz in a TissueLyser (Qiagen N.V., Venlo, The Netherlands) and were centrifuged for 15 min at 15000 g and 4 °C. Approximately 800 μL of the supernatant was transferred to an empty Eppendorf tube. The pellet was reextracted like described above, and subsequently, the supernatant from the second extraction was combined with the supernatant of the first extraction. Subsequently, 200 μL of the crude extract was transferred into LC–MS vials (1.5 mL ND9 bottle, Labsolute, Th. Geyer GmbH & Co. KG, Renningen, Germany) containing 800 μL extraction buffer.

### Chemical profiling of semi-polar metabolites

Reverse-phase liquid chromatography coupled to high resolution tandem mass spectrometry (LC-MS/MS) was used to profile extracts of different organs as described in (Anaia et al., 2025; Chiocchio et al., 2023). Briefly, semi-polar metabolites from the leaf extracts were separated employing a UPLC (Dionex UltiMate™ 3000, Thermo Fisher Scientific, Waltham, USA) equipped with a C18 analytical column (Acclaim TM RSLC 120; 2.1 × 150 mm, 2.2 µm particle size, 120 Å pore size). Briefly, the column temperature was kept at 40 °C. The mobile phase was composed of water (solvent A) and acetonitrile (solvent B), which both were acidified with formic acid (0.1% v/v). The chromatograph was set to a flow rate of 0.4 μL min^−1^ with a multi-step gradient (0 − 1 min 5%, 1 − 4 min 28%, 4 − 10 min 36%, 10 − 12 min 95%, 12 − 14 min 95%, 14 − 16 min 5%, 16 − 18 min 5%) for solvent B. The autosampler of the chromatograph was used to keep the leaf extracts at a constant temperature of 4 °C. The injection volume was set to 10 μL. For exact mass spectrometry conditions, see Anaia et al., 2025. The data was acquired in data-dependent acquisition (DDA) mode as described by Hazrati et al., 2022. Briefly, switch criteria for DDA MS/MS analysis were as follows: maximum three precursors per cycle with an absolute threshold for precursor selection of 5000 counts. The total cycle time was set to 2.4 seconds. Data were collected in positive ionization mode (ESI^+^), and the mass range was set to *m*/*z* 70–1500 Da. QTOF-MS parameters were set as follows: fragmentor voltage was set at 120 V, the capillary voltage was set at 4000 V, the skimmer voltage was set at 65 V, collision energy was set at 30 eV, drying gas temperature was set at 325 °C (8 L/min), nebulizing gas pressure was set at 40 psi, and sheath gas temperature and flow were set at 300 °C and 12 L/min, respectively. The resulting chromatograms were pre-processed using MetaboScape 5.0 (Bruker Daltonics GmbH, Bremen, Germany) as in Anaia et al., 2025.

### Chemometric metabolome mining

Mass spectra (.MGF) were exported from MetaboScape and analysed using SIRIUS (Böcker et al., 2009; Dührkop et al., 2019), which integrates a collection of tools, including CSI:FingerID/COSMIC (Dührkop et al., 2015; Hoffmann et al., 2022), ZODIAC (Ludwig et al., 2020) and CANOPUS (Dührkop et al., 2021). Using this pipeline, metabolomic features were categorized using the NPClassifier (Kim et al., 2021) and ClassyFire (Djoumbou Feunang et al., 2016) ontologies. A sum-normalized, log-transformed and auto-scaled peak intensity table containing all metabolomic features that were picked by MetaboScape was analysed using principal component analysis (PCA, Fig. 1) in *MetaboAnalystR 4.0* (Pang et al., 2021). Classified metabolomic features (all or a subset with NPClasses ‘steroids’, ‘triterpenoids’ and ‘pseudoalkaloids’) were used to perform non-metric multidimensional scaling (NMDS) using Bray-Curtis dissimilarity in *vegan*. Using this NMDS analysis, distinct clusters were assigned as *S. dulcamara* RDBE chemotypes (Fig. 2). Data of flowers and berries containing all picked features were subjected to Pareto scaling and the models (OPLS-DA) were developed and a shared-and-unique-structures (SUS) plot (Fig. 3A) was visualized using SIMCA v.18.0 (Sartorius Stedim Data Analytics AB, Umeå, Sweden). Supervised models were evaluated by the goodness-of -fit (R2x (cum) and R2y(cum)) and goodness-of-prediction (Q2(cum)), together with the parameters given by cross validation tests: permutation test (performed using 200 permutations) and Cross Validation Analysis of Variance (CV-ANOVA) (Table S1). The significance of the variables in each model was evaluated by their variable influence on projection (VIP) scores (Supporting material).

**Fig. 1:**
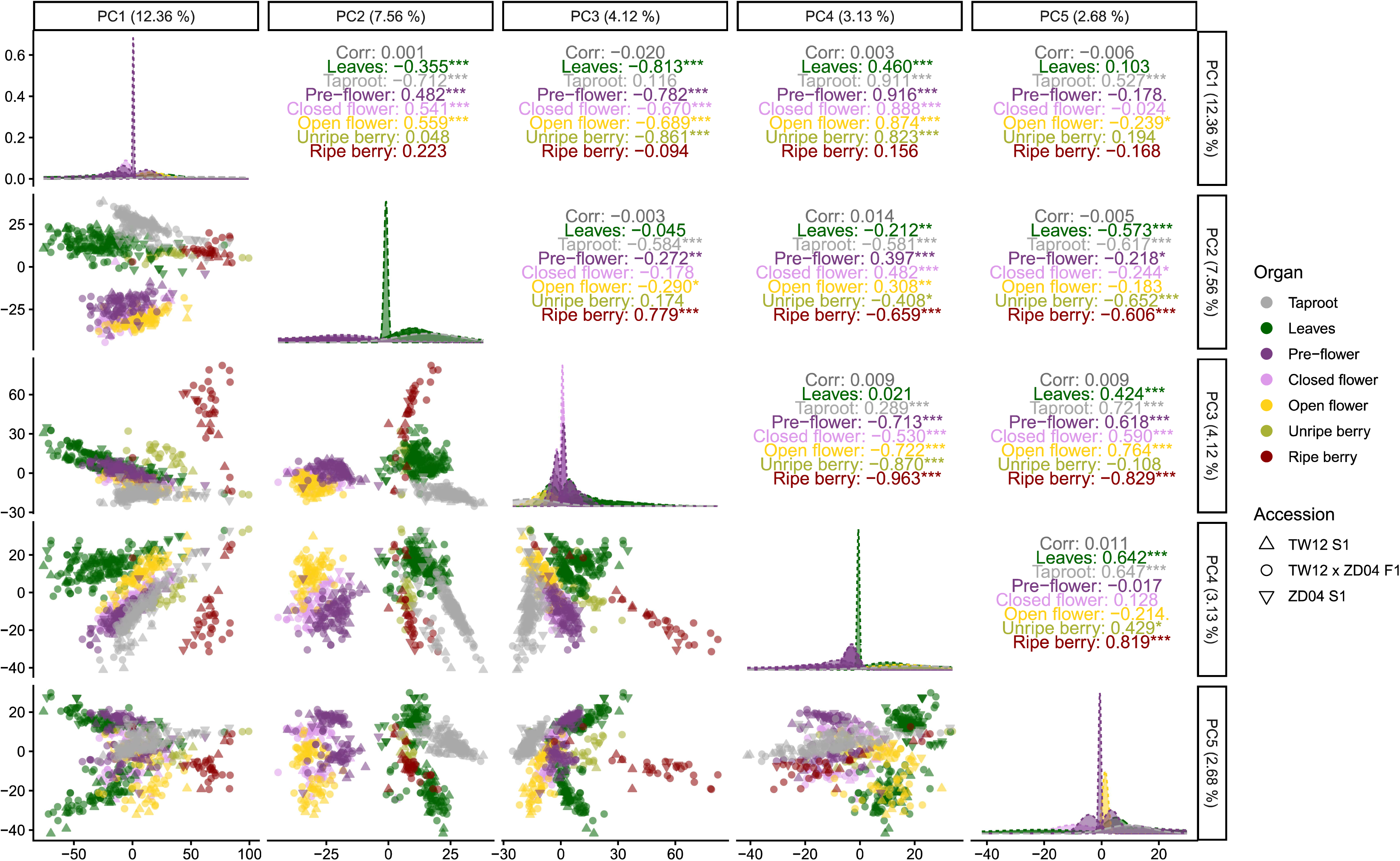
Generalized pairs plot of first five principal components (PC) from principal component analysis (PCA) of liquid-chromatography coupled with tandem mass spectrometry (LC-MS/MS) data. The PCA was based on all 6953 picked LC-MS/MS features for all injected samples. Each panel shows the bivariate relationship between two components, with distributions of individual components displayed along the diagonal. Node fill colour represents *Solanum dulcamara* organ (grey: roots; dark green: leaves; dark purple: pre-flowers, light purple: closed flowers; yellow: open flowers; olive green: pericarp and placental tissue [‘pulp’] of unripe berries and dark red: pulp of ripe berries). Shape represents genetic background (Triangle up: TW12 S_1_, diamond: TW12xZD04 F_1_ and triangle down: ZD04 S_1_). Tables depict Pearson correlation values and significance (asterisks: * p<0.05, ** p< 0.01, *** p<0.001).

**Fig. 2:**
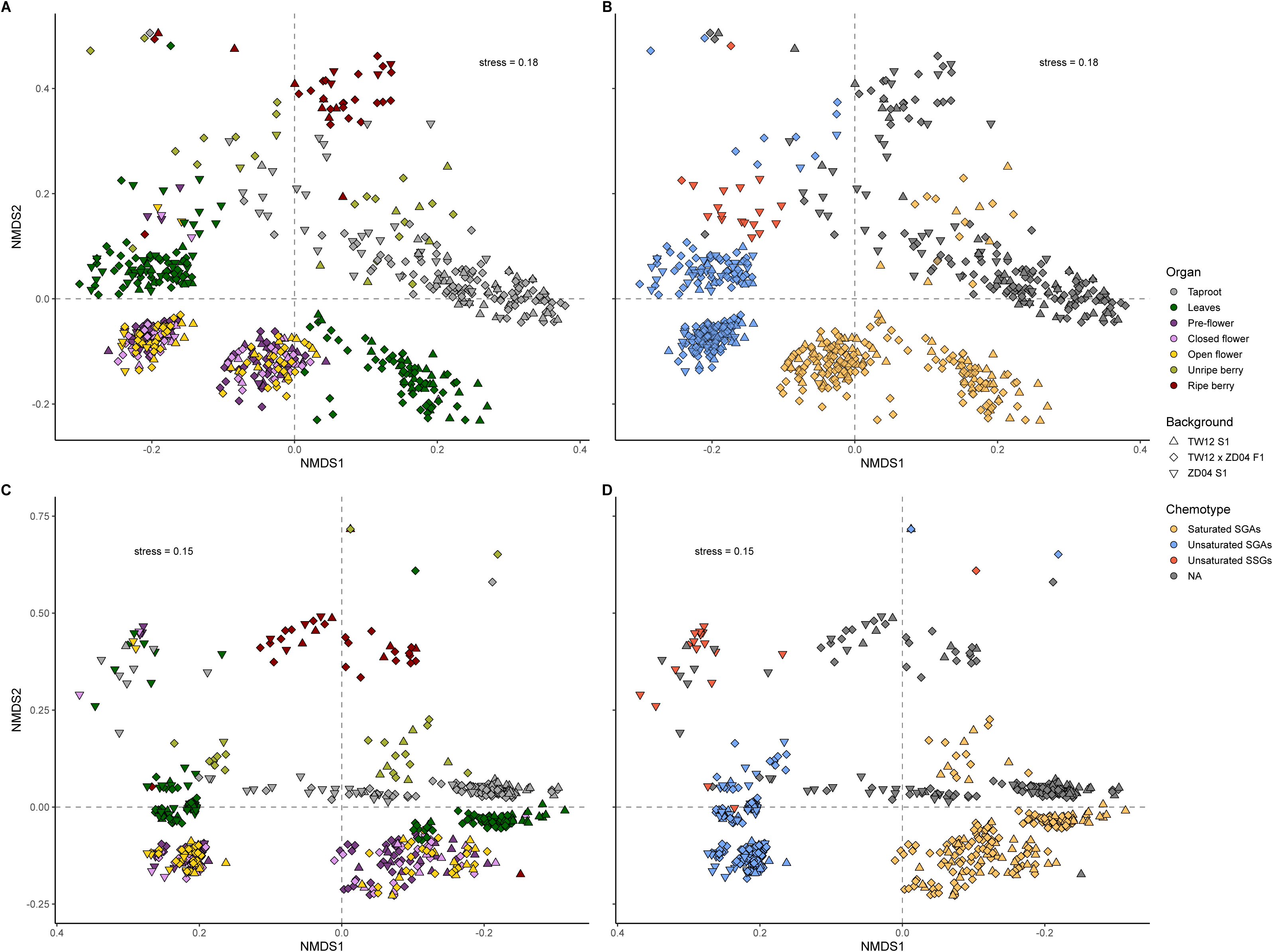
Non-metric multidimensional scaling (NMDS) using Bray-Curtis dissimilarity on classified LC-MS features of chemotypically segregating *Solanum dulcamara* progenies. For NMDS, visualization was performed either for 795 LC-MS/MS features with NPClassifier superclass annotations (*top panels;* A, B) or only retaining 634 features classified as ‘steroids’, ‘triterpenoids’ and ‘pseudoalkaloids’ (*bottom panels,* C, D). Node fill colour represents *Solanum dulcamara* organ (*left panels*, A, C; grey: roots; dark green: leaves; dark purple: flower buds, light purple: closed flowers; yellow: open flowers; olive green: pericarp and placental tissue [‘pulp’] of unripe berries and dark red: pulp of ripe berries) and chemotype (*right panels*, B, D; blue: predominantly unsaturated steroidal glycoalkaloids; yellow: predominantly saturated steroidal glycoalkaloids, red: predominantly unsaturated steroidal saponin glycosides). Shape represents genetic background (Triangle up: TW12 S_1_, diamond: TW12xZD04 F_1_ and triangle down: ZD04 S_1_).

**Fig. 3:**
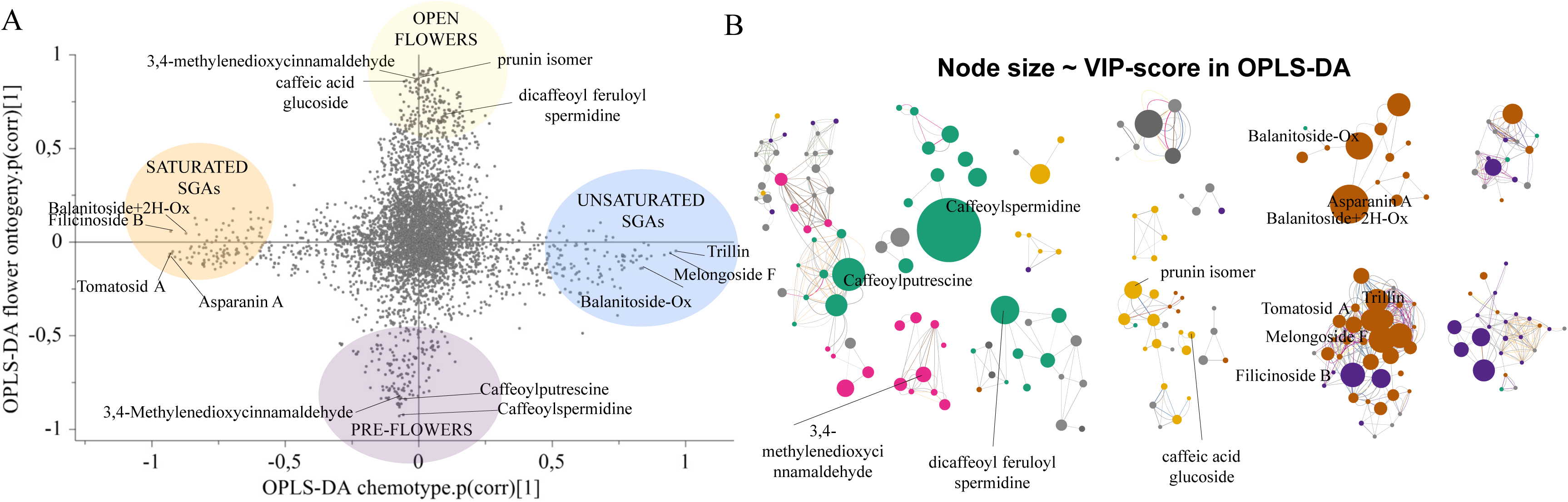
Chemometric molecular networking combining OPLS-DA with molecular networking. A) A) Shared and Unique Structures (SUS)-plot highlighting X variables (features) with the highest VIP-scores in the respective OPLS-DA models of flowers using flower chemotype (x-axis) and ontogeny (y-axis) as discriminant classes. B) Feature-based molecular networks combined with topic modelling. Node colours represent compound classes (brown: Steroids an d Triterpenoids; blue: Pseudoalkaloids (transaminated); green: Ornithine alkaloids; yellow: Flavonols; pink: Phenolic acids and Phenylpropanoids and dark grey: Coumarins). Node sizes are proportional to the VIP-scores derived from the OPLS-DA model discriminating classes based on flower development (y-axis, A).

Spectral library matching, feature-based molecular networking (FBMN, Nothias et al., 2020) and unsupervised topic modelling (MS2LDA, van der Hooft et al., 2016) were performed on the Global Natural Products Social Molecular Networking platform (GNPS, M. Wang, 2017; M. Wang et al., 2016). The outputs of aforementioned algorithms were integrated into a single molecular network using Cytoscape (Shannon et al., 2003). To prioritize networks for further exploration, node sizes in molecular networks were set in proportion to VIP-scores of the respective feature in the OPLS-DA models. For networks visualizing features from flowers and berries, the VIP scores from OPLS-DA models for flower and berry development were used, respectively. If not described otherwise, statistical models, post-hoc analyses and data visualizations were performed in R v. 4.3.3 using *lme4*, *emmeans* and *ggplot2*, respectively (Bates et al., 2015; R Core Team, 2024; Sievers et al., 2011; Wickham, 2016). Predicted SMILES were used to plot putative compounds using ACD/ChemSketch version 2020.1.2 (Advanced Chemistry Development, Inc., Toronto, Ontario, Canada).

## RESULTS

### Exploration of metabolic features using PCA

In order to analyse and identify metabolic differences among organs and the flower and fruit stages, we applied a series of progressive multivariate analyses. First, we performed PCA on all samples using all 6953 picked LC–MS/MS features (Fig. 1). Briefly, the first five principal components (PC) explained 29.9 % of the observed variance in the data (Fig. 1). Of the first five PCs, four could be associated with factors in our experimental design by comparing scores plot of PC1 with the other PCs (Fig. 1, left). Interestingly, PC1, PC2, P3 and PC5 correlated with variation in metabolites across ontogeny (PC1, not shown), among flowers (yellow & purple, PC2, PC3 & PC4), among ripe berries (red, PC2, PC3 & PC4) and likely across RDBE chemotypes (2 clouds of green dots for leaf samples, PC5, Fig. 1). Furthermore, in the scores plot between PC3 and PC4 (Fig. 1), green fruits (olive green) exhibit a clearer separation among chemotypes, consistent with the clustering observed in leaves (dark green), whereas red fruits show substantial overlap among chemotypes (Fig. 1).

### Leaf, flower and unripe fruit metabolomes show similar above-ground SG chemotypes

We hypothesized that part of the variation observed in the PCA was related to differences in SG profiles, in particular on PC5. Thus we followed up by non-metric multidimensional scaling (NMDS) using Bray-Curtis dissimilarity of 795 features with a NP classification (Fig. 2 A,B). Then, the NMDS analysis was repeated with a subset of 634 features that were annotated as ‘triterpenoids, steroids or pseudoalkaloids’ (Fig. 2C, D). Both analyses showed clear organ-specific (Fig. 2A) and chemotypic segregation (Fig. 2B) among samples of different *S. dulcamara* plants (Fig. 2). Ordination based on the annotated feature set showed that differences in leaf, flower and unripe berry chemotypes are detectable across metabolic profiles (Fig. 2B). NMDS restricted to computationally annotated SGs only (Fig. 2, bottom), sharpened this organ-specific (Fig. 2C) and chemotypic separation (Fig. 2D). Leaf, flower and unripe berry samples clustered consistently according to the two main chemotypes, U and S, independent of their lineage (Fig. 2A-D). This demonstrates that inherited SG chemotype is a major source of metabolomic differentiation in aboveground organs of *S. dulcamara*. Root and ripe fruit samples did not cluster in a similar way, confirming previous observations that root SG profiles of different leaf chemotypes converge (Anaia et al., 2025).

When considering SGs only, we found that flowers (yellow and purples) and leaves (dark green) and unripe fruits (olive green) co-segregated into two distinct clusters in the NMDS (Fig. 2C). This leads to the assumption that the *SdGAME25* expression polymorphism leading to either *U*- or *S*-type SGs in leaves also determines the chemotypes of flowers and unripe fruits (Fig. 2D, blue and yellow, respectively). Interestingly, samples associated with taproots (Fig. 2C, D, grey symbols) were mostly clustering with *S*-chemotype organs (Fig. 2D). This implies that taproots may contain relatively more *S*-type SGs, as observed before (Chiocchio et al., 2023). Furthermore, except for one leaf sample, leaf and flower samples, consisting of progeny from the ZD04-selfing (Fig. 2C, cloud of triangles in first quadrant, left) showed profiles indicative of a *U-*SSG chemotype (Fig. 2D, red symbols).

### Floral development and SG chemotype drive floral chemodiversity

Next, we analysed the subset of flower tissues using all picked metabolomic features to identify effects of flower development and chemotype. Orthogonal projections to latent structures discriminant analysis (OPLS-DA) separated pre-flowering buds (pre-flower) from open flowers, with strong predictive performance (Fig. S1, bottom; Table S1), indicating extensive metabolic remodelling towards anthesis. A second OPLS-DA model with all picked metabolomic features, based on the floral chemotypes classified using the NMDS, but excluding the unsaturated SSG chemotype (Fig. 1D, red symbols), also resolved distinct clusters (Fig. S1, top; Table S1), demonstrating that SG chemotypes remains evident in reproductive organs (Fig. S2A, 2B; Table S1).

To integrate both sources of variation, we generated a shared and unique structures (SUS) plot comparing the correlation between features and the first components of the ontogeny and chemotype OPLS-DA models (Fig. 3A). This analysis revealed metabolites specifically associated with transition from closed buds (pre-flower) to anthesis (open flowers; Fig. 3A, extremes on y-axis) and metabolites specifically associated with SG chemotype (Fig. 3A, extremes on x-axis). This means that developmental transitions across floral development reshape flower chemistry while retaining a chemotype-specific character. In addition to the metabolites specified in Fig. 3A, also some primary metabolites were found relevant in open flowers, namely amino acids and sugars (including monosaccharides and disaccharides) (Supporting information). Finally, we created molecular networks (Fig. 3B) in which the node size is proportional to the VIP scores extracted from the flower development OPLS-model (Fig. S1). Using this chemometric molecular networking approach, we were able to visually prioritize networks. Doing so, we identified networks containing nodes annotated as pseudoalkaloids, triterpenoids and steroids, flavonoids, polyamines, coumarins and phenylpropanoids as the most prominent (Fig. 3B). These networks were selected for further inspection.

### Steroidal saponins and glycoalkaloids jointly underpin chemotypic and ontogenetic variation

SGs were found to vary substantially during floral development (Fig. 3B). *S. dulcamara* flowers are predominantly characterised by the accumulation of SSGs, and to a lesser degree SGAs (Fig. 4A). In order to identify the driving factors of structural SG variation, node sizes were set proportional to the VIP-score from the OPLS-DA model for flower chemotype (Fig. 3A, x-axis).

**Fig. 4:**
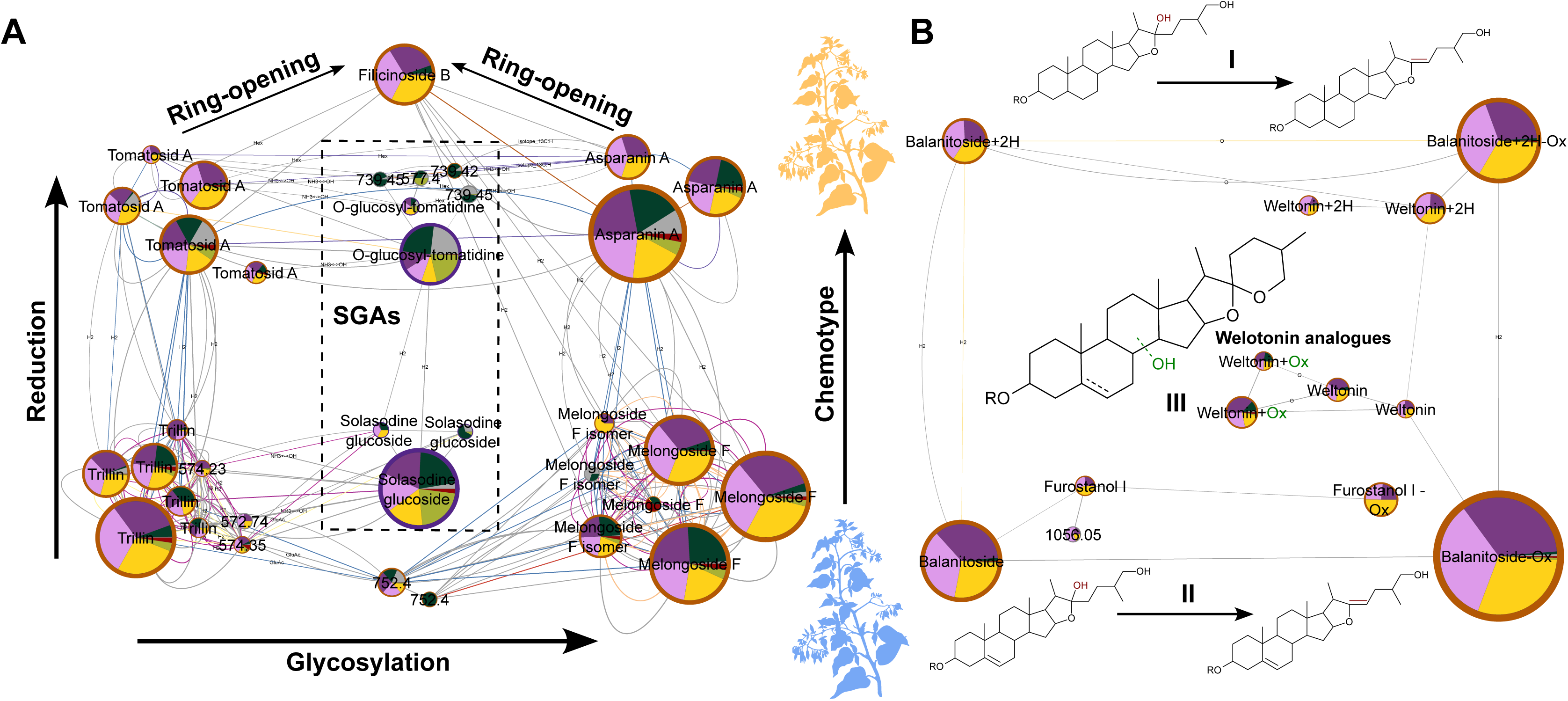
Biotransformations defining flower steroidal glycoside (SG) chemotypes revealed by feature-based molecular network combined with topic modelling. A) Steroidal saponin glycoside (SSG) biosynthesis (nodes arranged in perimeter) in flowers is associated with glycosylation, ring-opening and reduction, the latter leading to a similar polymorphism in SSGs as for the steroidal glycoalkaloid (SGAs, dashed rectangle) polymorphism previously observed in *Solanum dulcamara* leaves (blue plants: unsaturated SGAs; yellow plants: saturated SGAs). B) Flower-specific SSG biosynthesis (nodes arranged on perimeter) representing furostane steroids (**I-II**); nodes in the middle representing spirostane steroids **III**) defines flower chemotype identity. Node fill colour represents *Solanum dulcamara* organ (grey: roots; dark green: leaves; dark purple: pre-flowers, light purple: closed flowers; yellow: open flowers; olive green: pericarp and placental tissue [‘pulp’] of unripe berries and dark red: pulp of ripe berries). Node sizes are proportional to the VIP-scores derived from the OPLS-DA model discriminating classes based on flower chemotype (Fig. 3A, x-axis).

Factors driving structural SG variation are glycosylation (horizontal arrow, Fig. 4A) and ring-opening (towards filicinoside B isomers, Fig. 4A). However, the main chemotypic SG variation is characterised by differences in ring double bond equivalent between steroidal aglycons or chemical reduction (vertical arrow; Fig. 4A). Next to the flower chemotypic diversity, we also found that roots (grey) mostly contained U-SSG (Fig. 4A; top half), whereas S- and U-SGGs (green, olive green, red) were found more or less at the same frequencies in leaf and fruit samples.

Furthermore, we found another network that was associated with chemotypic SG variation in flowers only (Fig. 4B). This network consists of thirteen nodes, all classified in the NPC superclass ‘steroids’, except for one node, which was putatively misclassified as ‘ornithine alkaloid’ (Fig. 4B). Of the twelve nodes classified as steroids, six were classified as spirostanes and six as furostanes. Of the putative spirostanes, two nodes were annotated as molecular ions [M + H]+: 885.48 of weltonin (C_45_H_72_O_17_, MW: 884.48; VIP: 1.15 and 0.29; t_R_: 6.9 and 7.1 min., respectively), an unsaturated spirostane-type SSG with a triose glycoside moiety. Two other nodes (MW: 900.47), eluting at t_R_ 6.48 and 6.92 min., were connected to the weltonin nodes with a mass difference corresponding to the gain of an oxygen atom (ΔMW: +15.99), causing the weltonin+Ox nodes to elute earlier than their weltonin analogues (Fig. 4B). Noteworthy, is that the earlier eluting weltonin node, and the two weltonin+Ox nodes (VIP: 0.80 and 3.60; t_R_: 6.48 and 6.92 min., respectively), might be found in leaves next to flowers, which is not true for the later-eluting weltonin node. In general, the low VIP-scores suggest that spirostanol-type SSGs are common in both chemotypes, suggesting a conserved function for these spirostane-type SSGs in *S. dulcamara* organs.

Next to weltonin, another node is annotated as balanitoside (Fig. 4B, VIP: 14.12; t_R_: 6.9 min., MW: 1064.54, [M - H_2_O + H]^+:^ 1047.54), an unsaturated furostanol-type, bidesmodic SSG. Two nodes that elute later are putatively annotated as the saturated analogue of balanitoside (Fig. 4B, VIP: 3.46; t_R_: 7.0 min., balanitoside + 2H; MW: 1066.56, [M - H_2_O + H]^+:^ 1049.55) and saturated deoxybalanitoside (Fig. 4B, VIP: 7.68; t_R_: 7.2 min., balanitoside + 2H - Ox; MW: 1050.56; [M - H_2_O + H]^+:^ 1033.55), respectively. Furthermore, both balanitoside and its saturated analogue may lose an oxygen atom (ΔMW: −15.99), putatively annotated as deoxybalanitoside (balanitoside – Ox, VIP: 13.98 t_R_: 7.1 min., MW: 1030.54, [M + H] ^+:^ 1031.54), and balanitoside+2H-Ox (VIP: 7.68; t_R_: 7.21 min., MW: 1050.56, [M - H_2_O + H] ^+:^ 1031.54), respectively (Fig. 4B). Notably, the nodes associated with the furostanol-type SSGs are important drivers of SG chemotypic differentiation in *S. dulcamara* flowers (high VIP-scores, big nodes, Fig. 4).

### Conjugation of hydroxycinnamic acids to polyamines across floral development

To resolve metabolic changes underlying floral development (Fig. S1, bottom), we focused on metabolite classes enriched during the preflower**–**anthesis transition, in particular hydroxycinnamic acid (HCA) conjugates. We thereby found six clusters of HCAs and their respective conjugates as depicted in Fig. 5 A-F. Chemometric-molecular network analysis showed that pre-flowering stages contained simple monoconjugates formed by acylation of HCAs including caffeic and ferulic acid to spermidine, accumulating mainly caffeoylspermidine (VIP: 24.14; t_R_: 1.7 min.) and ferolylspermidine (VIP: 2.06; t_R_: 3.06 min.) (Fig. 5A, left). These aforementioned nodes formed a triad with another node annotated as caffeoylspermidine (VIP: 3.81; t_R_: 1.1 min.), suggesting that multiple amine groups can participate in HCA conjugation, giving raise to different isomers. Furthermore, another cluster contains two nodes that are 2 Da heavier than caffeoylspermidine (VIP: 2.63; t_R_: 1.09 min. and VIP: 1.74; t_R_: 1.38 min., respectively), suggesting that the double bond from the caffeic acid moiety may be (enzymatically) reduced (Fig. 5A). During floral development, these compounds are possible substrates that are further metabolised into increasingly substituted phenolamide analogues containing two or three HCA moieties conjugated to spermidine (Fig. 5A, right). Specifically, these compounds formed a network (Fig. 5A, right), consisting of sixteen nodes, of which seven nodes were categorized in the NP class of polyamines (Fig. 5A, right). Interestingly, all nodes except for two could be annotated as either di-*N*-acetylated (top cluster 5A, right) or tri-*N*-acetylated spermidine (bottom, Fig. 5A right). Some of the annotated spermidine-conjugates occur more than once in the network, suggesting the occurrence of different constitutional isomers. This can be explained by the fact that HCAs may be conjugated to spermidine at different amino groups (Table 2). Lastly, the annotated di-*N*-acetylated and tri-*N*-acetylated phenolamides occur at different proportions across floral development, suggesting that different spermidine-conjugates may have distinct biological functions. Alternatively, that the different nodes represent precursors in the biosynthetic pathway of these specific phenolamides. In our study, tri-*N*-acetylated phenolamides with a three identical HCAs conjugates were not detected in flower extracts, however two nodes putatively annotated as dicaffeoyl coumaryl spermidine and diferuloyl caffeoyl spermidine were predominantly detected in open flowers (yellow nodes, Fig. 3A), suggesting a specific role for these tri-*N*-acylated spermidine conjugates in open flowers or pollen. Similarly, the di*-N*-acylated spermidine conjugates dicaffeoyl spermidine and diferuloyl spermidine were predominantly detected in closed flowers, suggesting that these putative metabolites are the precursors for the tri-*N*-acylated spermidine conjugates predominantly detected in open flowers (Fig. 3A, Table 2).

**Fig. 5:**
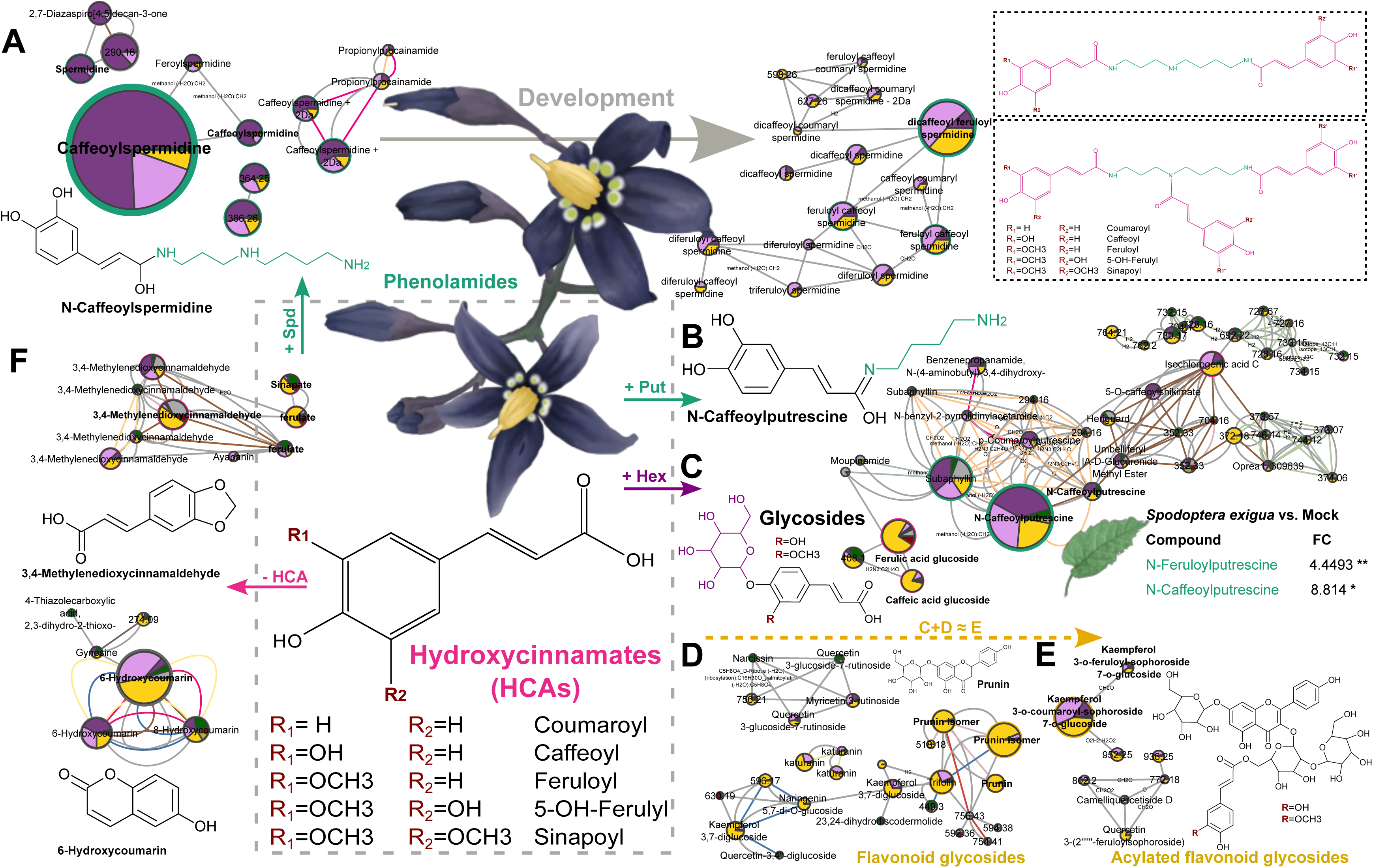
Acylation with hydroxycinnamates (HCAs, pink) is ubiquitous in *Solanum dulcamara* flowers. A) In pre-flowers spermidine-conjugates including caffeyolspermidine and feroylspermidine accumulate, which are transformed into di- and tri-acylated spermidine conjugates across floral ontogeny. B) Putrescine, is conjugated to caffeic acid raising caffeoylputrescine, which is found in all stages of floral ontogeny. C) Next to conjugation with polyamines, HCAs can be glycosylated. These glycosylated HCAs are mainly detected in open flowers, as is the case for D) flavonoid glycosides. Interestingly, E) acylated flavonoid glycosides, conjugates of C) HCA glycosides with D) flavonoid glycosides (yellow dashed arrow), are detected in *S. dulcamara* flowers. Next to conjugation to other compound classes, HCAs themselves are transformed into F) other compound classes such as cinnamaldehydes and coumarins in *S. dulcamara* flowers. Node sizes are proportional to the VIP-scores derived from the OPLS-DA model discriminating classes based on floral development (Fig. 2A, y-axis). Node fill colour represents *Solanum dulcamara* organ (grey: roots; dark green: leaves; dark purple: pre-flowers, light purple: closed flowers; yellow: open flowers; olive green: pericarp and placental tissue [‘pulp’] of unripe berries and dark red: pulp of ripe berries). Node circumference colours: blue: pseudoalkaloids, brown: steroids and triterpenoids.

**Table 2:** Putatively annotated spermidine-conjugates by manual propagation of annotations in a molecular network (Fig. 5A, right) in flowers of *Solanum dulcamara*. Bis: di-*N*-acylated spermidine. Tris: tri-*N*-acylated spermidine. Different amino groups are shown with N followed by an apostrophe.

| Spermidine-conjugate | Bis/Tris | N' | N'' | N''' |
| --- | --- | --- | --- | --- |
| Dicaffeoyl | Bis | Caffeoyl | H | Caffeoyl |
| Dicaffeoyl | Bis | Caffeoyl | Caffeoyl | H |
| Diferuloyl | Bis | Feruloyl | H | Feruloyl |
| Diferuloyl | Bis | Feruloyl | Feruloyl | H |
| Feruloyl caffeoyl | Bis | Feruloyl | H | Caffeoyl |
| Feruloyl caffeoyl | Bis | Feruloyl | Caffeoyl | H |
| Caffeoyl coumaryl | Bis | Caffeoyl | Coumaryl | H |
| Diferuloyl caffeoyl | Tris | Feruloyl | Feruloyl | Caffeoyl |
| Diferuloyl caffeoyl | Tris | Feruloyl | Caffeoyl | Feruloyl |
| Triferuloyl | Tris | Feruloyl | Feruloyl | Feruloyl |
| Dicaffeoyl feruloyl | Tris | Caffeoyl | Caffeoyl | Feruloyl |
| Dicaffeoyl coumaryl | Tris | Caffeoyl | Caffeoyl | Coumaryl |
| Dicaffeoyl coumaryl – 2H | Tris | Caffeoyl | Caffeoyl | Coumaryl – 2H |
| Feruloyl caffeoyl coumaryl | Tris | Feruloyl | Caffeoyl | Coumaryl |

In addition to spermidine, putrescine is another major floral polyamine that is acylated with different HCAs, which drive metabolic separation between pre-flowers and open flowers. (Fig. 3B). These HCA – putrescine conjugates, such as coumarylputrescine (VIP: 2.63; t_R_: 1.09 min.) and caffeoylputrescine (VIP: 11.62; t_R_: 3.06 min. and VIP: 2.18; t_R_: 2.4 min., respectively), are predominantly detected in extracts of *S. dulcamara* flowers, though trace-levels are present in leaves as well (Fig. 3B, green sectors in nodes). Interestingly, when *S. dulcamara* leaves are challenged with herbivory posed by *S. exigua* larvae for 24 hours, the levels of caffeoylputrescine (fold-change: 4.4, *P <* 0.05), together with feroylputrescine (fold chance: 8.8, *P <* 0.01; Fig. 3B, Table 3) both significantly increase. This suggests that reproductive organs constitutively deploy HCA – putrescine conjugates otherwise associated with inducible chemical defence responses in leaves.

**Table 3:** LC-MS features and annotated fragments induced after 24 hours *Spodoptera exigua* feeding. Features are shown as retention time in seconds followed by *m/z*-value, separated by a slash.

| Feature or annotated fragment | Relative fold change<br>(*Spodoptera* vs. Mock) | Adjusted<br>*P*-value |
| --- | --- | --- |
| 413.74/313.13113 | 48.437 | 0.0067565 |
| **6-Hydroxy-7-methoxycoumarin** (Fig. 5F) | 28.52 | 0.0067565 |
| 698.55/672.27924 | 10.138 | 0.0067565 |
| 243.1/192.04241 | 5.998 | 0.0067565 |
| Phenylacetaldehyde | 4.5735 | 0.0067565 |
| **Feruloyl putrescine** (Fig. 5B) | 4.4493 | 0.0067565 |
| 87.2/137.08386 | 4.0496 | 0.0067565 |
| 330.86/692.26885 | 7.246 | 0.0072069 |
| 57.04/295.05851 | 4.9072 | 0.0072069 |
| 57.75/257.1033 | 4.7857 | 0.0072069 |
| 243.87/334.11617 | 4.337 | 0.0072069 |
| **Caffeoyl putrescin** (Fig. 5B) | 8.814 | 0.022257 |

### Glycosylation of HCAs and further acylation with flavonoid glycosides in open flowers

HCAs are also present as glycosides, which are particularly abundant in open flowers (Fig. 5C). Conjugation to glycosides increase polarity and thereby vacuolar storage capacity and transport potential, thereby representing another route of HCA diversification during anthesis. Next to these HCA glycosides, flavonoid glycosides including putatively annotated prunin isomers (3 nodes; VIPs: 5.52, 3.16 and 2.17 at t_R_: 6.17, 5.59, and 5.02 min., respectively), diglucosides of kaempferol (2 nodes; VIPs: 2.17 and 2.13 at t_R_: 4.09 and 4.72 min.,) and naringenin (VIP: 0.99; t_R_: 4.71 min.), and rutinosides of myrcetin (VIP: 1.29 ; t_R_: min.) and quercetin 3-glucoside (VIP: 0.80; t_R_: 4.47 min.) mainly accumulate in open flowers (Fig. 5D). Flavonoid glycosides can be further acylated in *S. dulcamara* flowers, giving rise to acylated flavonoid glycosides, which contain an HCA moiety (Fig. 3E). For example, two nodes putatively annotated as kaempferol 3-O-coumaroyl-sophoroside 7-O-glucoside (VIP: 6.42; t_R_: 4.43 min.) and kaempferol 3-O-ferulyl-sophoroside 7-O-glucoside (VIP: 0.75; t_R_: 4.54 min.) contain coumaric and ferulic acid moieties conjugated at the 3-hydroxyl position of kaempferol, respectively (Fig. 5E). Finally, part of the HCA pool appeared to be redirected toward formation of cinnamaldehydes and coumarins (Fig. 5F), instead of being conjugated with glycosides. This suggests that anthesis involves fluxes into multiple phenylpropanoid end-products rather than accumulation of a single metabolite class in *S. dulcamara*. Noteworthy are two nodes annotated as 6-hydroxcoumarin and five nodes annotated as 3,4-methylenedioxycinnamaldehyde (Fig. 5F). The nodes with the highest VIP-scores for 6-hydroxycoumarin (VIP: 2.83; t_R_: 4.06) and 3,4-methylenedioxycinnamaldehyde (VIP: 0.71; t_R_: 4.32) are associated with open flowers (Fig. 5F). Interestingly, a 7-methoxylated analogue of 6-hydroxycoumarin accumulates (fold-change: 28.52, *P <* 0.01) after *S. exigua* herbivory (Table 3). Collectively, these data identify floral development as a period of intense phytochemical rearrangement and diversification centred around HCA accumulation and glycosylation as well as acylation of polyamines and flavonoid glycosides.

### N- and O-acetylation of SGAs during fruit ripening

Using LC-MS data from berry extracts, we also investigated the metabolic shifts associated with fruit ripening. OPLS-DA clearly separated extracts from unripe and ripe berry pulp, demonstrating that fruit ripening development is accompanied by major shifts in metabolic profiles (Fig. S2A). Whereas diversification of phenylpropanoid-derived PSMs dominated across floral development, fruit ripening is strongly characterised by acetylation of *S. dulcamara* steroidal glycoalkaloids (Fig. S2B). Fragment *m/z* fragments indicating SGAs, such as solasonine, solamargine and soladulcin A, are strongly associated with extracts from all organs, including green berry pulp, but not with red berry pulp (Fig. 6A). Specifically, two solasodine (VIP: 14.15 and 7.57 at t_R_: 5.68 and 5.81 min., respectively) and two soladulcidine (VIP: 7.07 and 3.57 at t_R_: 6.02 and 6.3 min., respectively) isomers are converted into six 28N-acetylspirosolan-3-ol (VIP: 1.88, 1.57, 1.30, 0.99, 0.77, and 0.66; t_R_: 9.10, 8.59, 7.03, 8.9, 7.73 and 11.76 min.) and seven 28N-acetylspirosolan-5-en-3-ol (VIP: 4.65, 4.15, 2.66, 1.13, 0.62, 0.55 and 0.48 at t_R_: 8.42, 6.95, 8.72, 7.86, 11.05, 9.46 and 7.48 min.) isomers, respectively (Fig. 6A). This suggests that during fruit ripening SGA conversion mechanisms target the steroidal aglycone. This is further corroborated by the fact that of two nodes, putatively annotated as N-acetylated analogues of SGAs, such as 28N-acetylsolasonine (MW: 909.508 Da; VIP: 10.64; t_R_: 8.72 min.), one node is associated with berry pulp *in general*, whereas the other node, 28N-acetylsolasonine (MW: 909.508 Da; VIP: 2.76; t_R_: 11.79 min.) is only associated with red berry pulp (Fig. 6B). Three nodes were annotated as 28N-acetylsolamargine (MW: 925.503 Da; VIP: 3.77, 1.27 and 0.36; at t_R_: 8.42, 11.75 and 11.06 min.), of which the two latter nodes are associated *only* with red berry pulp (Fig. 6B). This indicates that the biosynthetic pathway towards acetylation begins prior to full berry ripening, but becomes increasingly prominent during maturation (Fig. 6B). After hydroxylation, O-acetylation of steroidal aglycons can occur, potentially producing a larger number of modified aglycon derivatives compared with the N-acetylation pathway (Fig. 6B). These reactions substantially increase structural diversity within the berry steroidal metabolome, as witnessed by two nodes annotated as O-acetylsolasonine (MW: 941.50 Da; VIP: 0.48 and 0.15 at t_R_: 6.96 and 7.83 min.) and one node as O-acetylsoladulcidine (MW: 943.51 Da; VIP: 2.76; t_R_: 11.79 min., Fig. 6B). Furthermore, several O-acetylated SGAs are associated with oxidative deamination pathways leading toward SSGs such as tomatonin (MW: 902.49; VIP: 1.08; t_R_: 5.33 min.) and tomatonin - 2 Da (MW: 900.47; VIP: 1.54; t_R_: 5.33 min., Fig. 6C). Such transitions are consistent with detoxification processes, as removal or modification of nitrogen-containing functional groups may reduce biological activity (Zhao et al., 2021). Lastly, a final set of predicted berry-associated steroidal aglycons suggested partial steroid degradation products, many of which were associated with acetylation as well (Fig. 6D). This pattern implies that N- and O-acetylation accompanies multiple stages of steroidal metabolite turnover, from early modification of canonical SGAs to downstream detoxification and degradation products.

**Fig. 6:**
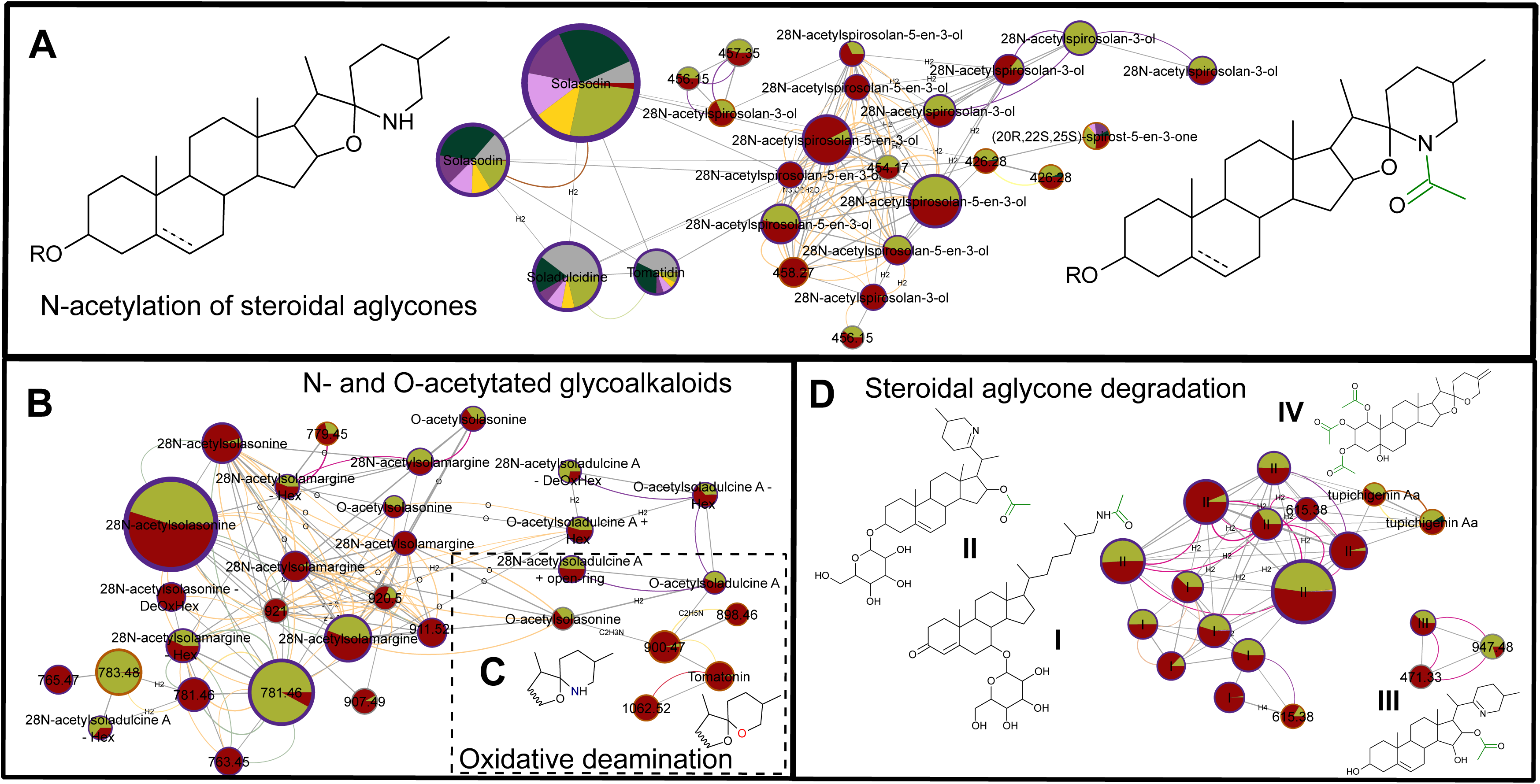
Biotransformation of steroidal glycoalkaloids (SGAs) by N- and O-acetylation across fruit ripening revealed by feature-based molecular network combined with topic modelling. A) N-acetylation of steroidal aglycones (SAs). The common *Solanum dulcamara* SGAs (top-left) are detected in all tested organs (4 top-left multi-coloured nodes) except in pulp of ripe fruits. However, their ‘detoxification products are only detected in unripe (olive green nodes) and ripe (red nodes) berry pulp. B) Both N- and O-acetylated steroidal glycoalkaloids (SGAs) are detected, indicating two distinct SGA detoxification mechanisms. C) Subsequently, O-acetylated steroidal glycosides (e.g., O-acetylsolasonine and O-acetylsoladulcine A) can be biotransformed into steroidal saponin glycosides by oxidative deamination via an open-ring intermediate. D) Steroidal aglycone degradation (e.g., ring-opening, conversion from secondary to tertiary amine, acetylation of hydroxyl-groups) is associated with both N- and O-acetylation. Chemical structures (**I-IV**) shown are *in silico* predictions. Node sizes are proportional to the VIP-scores derived from the OPLS-DA model discriminating classes based on berry development (Fig. S2). Node fill colour represents *Solanum dulcamara* organ (grey: roots; dark green: leaves; dark purple: pre-flowers, light purple: closed flowers; yellow: open flowers; olive green: pericarp and placental tissue [‘pulp’] of unripe berries and dark red: pulp of ripe berries).

## DISCUSSION

Poisonous plants are commonly characterised by their dominant phytochemicals, yet toxicity-related metabolites often varies substantially among individuals, organs and across development. Increasing evidence shows that plant specialised metabolism is tightly regulated in space and time, reflecting dynamic ecological demands (Barton & Koricheva, 2010; Moore et al., 2014). Our results, show strong metabolic separation among leaves, flowers, and fruits as well as among developmental stages within flowers and fruits, supporting a more eco-physiological view of plant chemical defences. Developmental control of toxicity is likely adaptive, because reproductive organs experience contrasting selection pressures over time. Flowers must remain functional and attractive to pollinators while resisting antagonists, whereas fruits frequently shift from defence during immature stages to attraction of seed dispersers during ripening (Cipollini & Levey, 1997a; Tewksbury & Nabhan, 2001). The extensive chemical restructuring observed here in *S. dulcamara* is consistent with these life-stage specific ecological trade-offs.

Previous work in *Solanum dulcamara* and related species has demonstrated heritable intraspecific variation in steroidal glycosides and other defence metabolites (Anaia et al., 2025; Calf et al., 2018; Chiocchio et al., 2023; Eich, 2008; Köthe & Willuhn, 1983; Sander & Willuhn, 1961; Willuhn, 1966, 1967, 1968, 1969; Willuhn & Kun–anake, 1970). Our finding that floral tissues also segregate by SG chemotype indicates that these polymorphisms are expressed systemically across aboveground plant tissues. Previously, we associated *SdGAME25* expression with SG organ- and development-specific chemodiversity (Anaia et al., 2025; Calf, 2019). Our current results make it likely that changes in *SdGAME25* expression are also responsible for determining floral chemotype. Whole-plant expression of SG chemotype may have important ecological consequences, because floral chemistry can affect florivores, pollen feeders, microbial communities, and pollinators (Adler, 2000; Stevenson et al., 2017). Specialised metabolites in nectar, pollen, and floral tissues are increasingly recognised as determinants of plant–pollinator interactions and reproductive success (Adler, 2000; Manson et al., 2013). Thus, inherited variation in poisonous SGs may influence multiple trophic interactions associated with plants simultaneously.

The transition from preflower buds to open flowers was characterised by pronounced enrichment of hydroxycinnamic acid (HCA) conjugates, flavonoid glycosides, and polyamine derivatives. Such metabolites are widely associated with reproductive tissues in angiosperms and often peak during anthesis (Fellenberg & Vogt, 2015). In our study, progressive accumulation of acylated spermidine conjugates was particularly notable. Hydroxycinnamoyl spermidines occur commonly in pollen coats and anthers, where they have been implicated in pollen wall development, UV protection, and defence against microbial attack (Elejalde-Palmett et al., 2015). In *Arabidopsis thaliana*, trihydroxycinnamoyl spermidine derivatives accumulate in the pollen coat and were found to be synthetized by a spermidine hydroxycinnamoyltransferase. In apple trees, N1,N5,N10-tricoumaroyl spermidine and N1,N5-dicoumaroyl-N10-caffeoyl spermidine accumulate specifically in pollen grain coat (Elejalde-Palmett et al., 2015). Their enrichment in open flowers therefore suggests functional importance during maximum reproductive exposure. In pre-flowering buds, caffeoylputrescine was also abundant and driving separation between preflowers and open flowers. In several plant species, caffeoylputrescine and related phenolamides are induced by herbivory or jasmonate signalling, where they contribute to anti-herbivore defence (Calf et al., 2020; Kaur et al., 2010; Onkokesung et al., 2012). Their constitutive presence in flowers may indicate pre-emptive protection of high-value tissues, such as emerging reproductive organs. Flavonoid glycosides and acylated flavonoids are generally linked to floral pigmentation, UV protection, antioxidant capacity, and pollinator attraction (Agati et al., 2012). Their coordinated accumulation indicates that flower opening is accompanied by a broad functional reprogramming of specialised metabolism. In *Arabidopsis thaliana*, trihydroxycinnamoyl spermidine derivatives accumulate in the pollen coat and were found to be synthetized by a spermidine hydroxycinnamoyltransferase. In apple trees, N1,N5,N10-tricoumaroyl spermidine and N1,N5-dicoumaroyl-N10-caffeoyl spermidine accumulate specifically in pollen grain coat (Elejalde-Palmett et al., 2015). The coexistence of chemotypic and ontogenetic signals further suggests that constitutive genetic differences are layered upon developmental regulation, generating chemically distinct floral phenotypes at each developmental stage.

In contrast to the flowers, fruit maturation was dominated by the detoxification of SGAs. Unripe fruits contained mainly common SGAs, whereas ripe berry pulp more often contained N- and O-acetylated SGAs. In cultivated tomato, toxic glycoalkaloids such as α-tomatine decline during fruit ripening and are converted into less toxic esculeosides through hydroxylation, glycosylation, and related transformations (Iijima et al., 2009; Martínez-Rivas & Fernie, 2024; Nohara et al., 2010; Yamanaka et al., 2009; Zhu et al., 2022). Similar ontogenetic declines in glycoalkaloid toxicity have been documented in other fleshy-fruited species, where immature fruits require defence, but ripe fruits benefit from vertebrate dispersers (Cipollini & Levey, 1997a; Whitehead & Bowers, 2013). Acetylation may further reduce biological activity by altering polarity, membrane affinity, or accessibility of reactive functional groups. For example, it was found that bioactivity of SGAs decreases upon E-ring opening or acetylation of the amide in the F-ring (Friedman, 2006, 2015; Zhao et al., 2021). Although the specific bioactivities of the N- and O-acetylated SGA derivatives detected here remain to be tested, our results strongly support a conditional defence model in which berry chemistry shifts from deterrence to dispersal compatibility during seed maturation.

While untargeted metabolomics and chemometric molecular networking provide strong evidence for developmental restructuring, structural confirmation of several predicted compounds will require targeted isolation and NMR-based characterisation. Functional assays using herbivores, pathogens, and frugivores are also needed to determine how acetylation and related modifications alter realised toxicity. Future work should additionally investigate the enzymes underlying these transitions, particularly BAHD acyltransferases (Sonawane et al., 2023), glycosyltransferases, and dioxygenases (Cárdenas et al., 2019) known to diversify defence chemistry in Solanaceae (Moghe & Last, 2015; Schilmiller et al., 2015).

Despite major changes in overall metabolic profiles during flower and fruit development, steroidal glycosides remained dominant contributors to the chemical profiles of all organs. This reinforces their central role in defence across Solanaceae (Friedman, 2006; Milner et al., 2011). Importantly, toxicity and/or deterrence is unlikely to depend on total glycoside concentration alone. Small structural differences in aglycone saturation, hydroxylation state, glycosylation pattern, or acylation can substantially alter membrane disruption, bitterness, digestibility, and antimicrobial activity (Cárdenas et al., 2015; Friedman, 2015). Consequently, studies of poisonous plants must consider both concentration and compositional turnover as well as disparity (Petrén et al., 2024).

## Conclusion

Our findings suggest that poisonous chemistry in *S. dulcamara* is shaped by multiple selective agents acting across reproductive development. Floral metabolites likely reflect simultaneous pressures from florivores, pathogens, and pollinators, whereas changes in fruit metabolic profiles reflect opposing pressures from seed predators and dispersers. These ontogenetically orchestrated changes add to the overall chemodiversity of the leaf chemotypes in *S. dulcamara* (Calf et al., 2018; Willuhn, 1966). Chemotypic polymorphisms may be maintained when alternative chemical phenotypes confer context-dependent fitness benefits, as predicted under balancing selection for defence traits (Christie & McNickle, 2023; Garrido et al., 2016; Goldberg et al., 2020; Wittmann & Bräutigam, 2024). Developmental reprogramming may further expand adaptive potential by allowing a single genotype to express different toxic phenotypes through time. More broadly, these results illustrate how poisonous plants can reconcile defence with reproduction through temporally dynamic deployment of plant metabolites.

## Acknowledgement

We thank Alvin Q. Barth for support in the greenhouse. We thank Dr. Stefanie Döll for expert support during data acquisition. We thank Arjin Koç, Dr. Ming Zeng and Hanna Ziegler for practical assistance. Special thanks go to Dr. Janny L. Peters, Radboud Institute for Biological and Environmental Sciences, Radboud University, Nijmegen, the Netherlands, for hosting a guest stay of RAA, which included access to facilities and discussions on the research presented in this paper.

## Author contributions

RAA and NMvD conceived and designed this study. NMvD made crosses to obtain the studied plant accessions and obtained funding. RAA conducted the experiments, analysed the resulting data, prepared figures and wrote a draft. IC built OPLS-DA models, prepared figures and provided input for the draft. All authors agreed with submission of current manuscript to the journal.

## CRediT statement

RAA – Conceptualization, Data Curation, Formal Analysis, Investigation, Methodology, Project administration, Validation, Visualization, Writing – original draft, Writing – review & editing.

IC – Data curation, Formal analysis, Resources, Validation, Visualization, Writing – review & editing

NvD – Conceptualization, Funding acquisition, Methodology, Project administration, Resources, Supervision, Writing – original draft, Writing – review & editing.

## Funding information

RAA and NMvD gratefully acknowledge the German Research Foundation (DFG) for funding to the Research Group ChemDiv (DFG-FOR 3000/1, P4 DA 1201/10-1, 415496540) and iDiv (DFG-FZT 118, 202548816). RAA was funded by an ERAMSUS+ grant from the European Union for the guest stay at Radboud University.

## Data availability statement

Raw LC-MS data that are presented in this study are made publicly available through MassIVE (MSV000100619) upon publication.

---

**Table S1.**
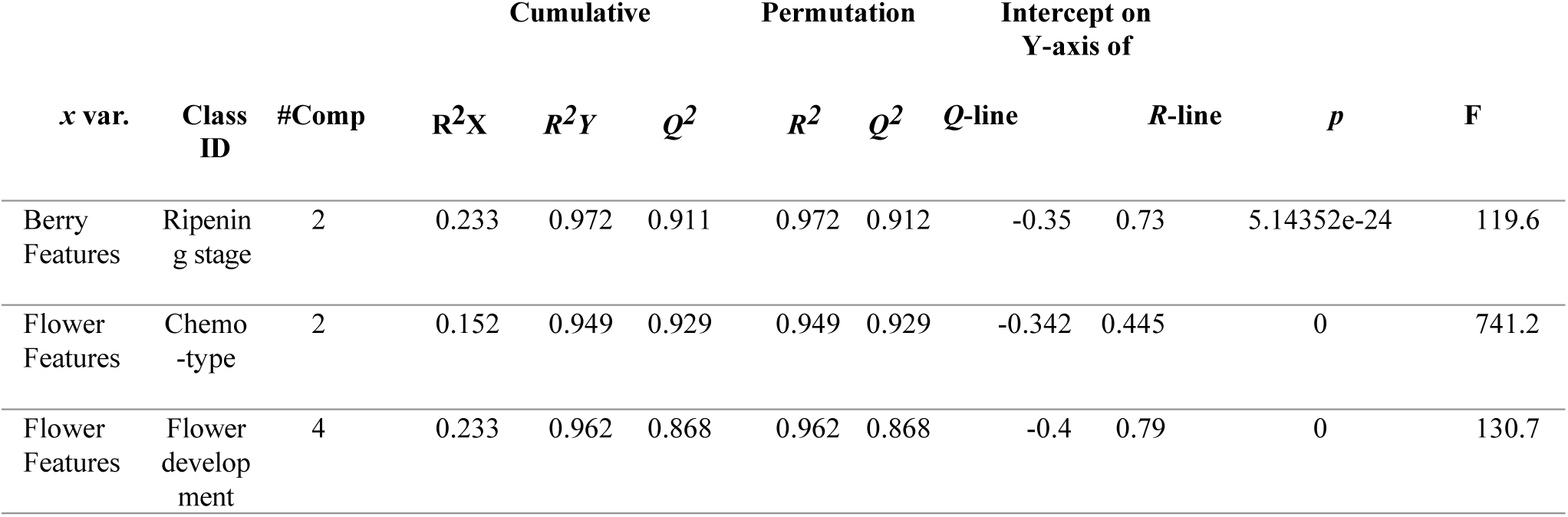
OPLS-DA models developed in this study with their parameters (R^2^X cum) and Q^2^(cum), and the parameters for the validation obtained by permutation test (200 permutations) and CV-ANOVA.

| x var. | Class ID | #Comp | Cumulative |  |  | Permutation |  | Intercept on Y-axis of |  | p | F |
| --- | --- | --- | --- | --- | --- | --- | --- | --- | --- | --- | --- |
|  |  |  | R<sup>2</sup>X | R<sup>2</sup>Y | Q<sup>2</sup> | R<sup>2</sup> | Q<sup>2</sup> | Q-line | R-line |  |  |
| Berry Features | Ripening stage | 2 | 0.233 | 0.972 | 0.911 | 0.972 | 0.912 | -0.35 | 0.73 | 5.14352e-24 | 119.6 |
| Flower Features | Chemotype | 2 | 0.152 | 0.949 | 0.929 | 0.949 | 0.929 | -0.342 | 0.445 | 0 | 741.2 |
| Flower Features | Flower development | 4 | 0.233 | 0.962 | 0.868 | 0.962 | 0.868 | -0.4 | 0.79 | 0 | 130.7 |

## FIGURE CAPTIONS

**Fig. S1: OPLS-DA models of flowers using flower chemotype (A, B) and development (C, D) as discriminant classes.** Scores plot (left panels) of pre- (left, green) and closed (red, right) flowers of *Solanum dulcamara* plants from different genetic backgrounds (triangle up: TW12 S1, triangle down: ZD04 S1, diamond: TW12 x ZD04 F1). S-plots highlighting LC-MS(/MS) features (right panels) associated with pre- (bottom-left quadrant) and open- (top-right quadrant) flowers. The dimension of loadings (features) is proportional to their VIP value.

**Fig. S2: OPLS-DA models of berries using berry development as discriminant class.** A) Scores plot of unripe (left, green) and ripe (red, right) berry pulp of *Solanum dulcamara* plants from different genetic backgrounds (triangle up: TW12 S1, triangle down: ZD04 S1, diamond: TW12 x ZD04 F1). S-plots highlighting LC-MS(/MS) features (right panels) associated with pre- (bottom-left quadrant) and open- (top-right quadrant) flowers. The dimension of loadings (features) is proportional to their VIP value. The dimension of loadings (features) is proportional to their VIP value. 1: solamargine isomer [M+H]+, 2: α-soladulcine isomer [M-Hex-Pent]+, 3: solasonine isomer [M+H]+, 4: solasonine [M-solatriose+H]+, 5: solamargine isomer [M+H]+, 6: 28N-acetylsolasonine isomer - 4 Da [M+H]+, 7: amino acid I (M = 123.99506, t_R_: 40.31 s), 8: steroidal alkaloid III [M+H]+ (M = 889.47877, t_R_: 365.64 s), 9: soladulcine A isomer [M+H]+; 10: 28N-acetylsolasonine isomer [M+H]+, 13: steroidal alkaloid II (M = 915.48258, t_R_: 259.53 s), 14: unknown compound (M = 172.07135, t_R_: 177.69 s), 15: unknown compound (M = 202.04547, t_R_: 58.1 s), 16: steroidal alkaloid I (M = 899.48743, t_R_: 264.02 s)

## Appendix S1

### Caterpillar feeding experiment

*Solanum dulcamara* plants of genotype “TW12” were grown from stem cuttings of at least 3 centimetres consisting of multiple nodes. Accession “TW”, of which “genotype TW12” is a previously studied individual, was originally collected as seeds from a population located in a wetland on Texel, an island of the Netherlands (53°07’17.5"N; 4°47’13.6"E) and germplasm is stored at the Centre for Genetic Resources (Wageningen, The Netherlands; Radboud University Genebank Accession nr. B24750045). Stems with multiple nodes were cut into segments of approximately 3 cm and were planted in plastic pots (11 x 11 x 12 cm) (Lamprecht-Verpackungen GmbH, Göttingen, Germany) containing a mixture of 50% sand, 50% ‘Floradur® B pot clay medium’ potting substrate (Floragard Vertriebs-GmbH, Oldenburg, Germany) supplemented with 4 g/L ‘Osmocote® Pro’ fertilizer (ICL Specialty Fertilizers, Nordhorn, Germany). Plants were grown in a greenhouse with ample water supply to stimulate adventitious root formation. Plants were grown under greenhouse conditions (24 °C at 16 h day and 17°C at 8 h night, mean temperature of 22°C at 70% relative humidity) and were watered every other day. All experimental units that were used for experimentation were 7-week old, well-watered plants and were selected based on phenotypic appearance and overall vigour. Beet armyworm [*Spodoptera exigua* (Lepidoptera, Noctuidae)] eggs were commercially sourced (Entocare C.V., Wageningen, The Netherlands) and insects were reared for many generations in a climate-controlled (25 °C at 45% relative humidity, 12 h/12 h day-night cycle) growth chamber (E-36L2c, Percival Scientific Inc, Perry, IA, USA). Eggs were hatched and insects were reared in vented plastic boxes (11.5 x 11.5 x 6 cm) and were fed daily with artificial diet from neonate until pupae stage. Moths were fed using a piece of cotton saturated in a 20% organic honey solution, and were kept in insect cages (38 x 38 x 60 cm; Entomologie-Speciaalzaak Vermandel V.O.F., Hulst, The Netherlands) with filter paper as substrate for oviposition. Fifth and sixth instar larvae were used for experimentation. Briefly, plants of accession “TW12” were grown as described above and were supported on bamboo sticks. 11 seven-week old plants were selected based on homogenic phenotypic appearance and vigour, and were assigned to a mock- (n=8) or herbivory- (n=9) treatments. The 16^th^ leaf counting from the first fully expanded leaf at the apex of the plants was introduced into a clip-cage and either challenged or not with 7-8 sixth instar *S. exigua* larvae for 24 hours. Top and bottom halves of clip cages were held together with rubber bands and were clipped onto the bamboo sticks using steel wires. To ensure that enough leaf material remained for harvesting in the herbivory treated plants, clip cages were put onto the leaf apex, excluding the leaf base. Treated leaves were harvested using scissors 24 hours after start of the treatment, and were put into a 15 mL Falcon tube, which was subsequently flash-frozen in liquid nitrogen. Flash-frozen samples were stored at −80 °C until further processing. Furthermore, mock- and treated-plants were physically separated by growing them on different tables within the greenhouse compartment.

## Notes

### Competing Interest Statement

The authors have declared no competing interest.

