## Supplementary figures and images for "Bittersweet dynamics from flowers to fruits: chemometric molecular networking reveals metabolic changes towards reduced toxicity over ontogeny in above-ground *Solanum dulcamara* chemotypes"

### Fig. S1

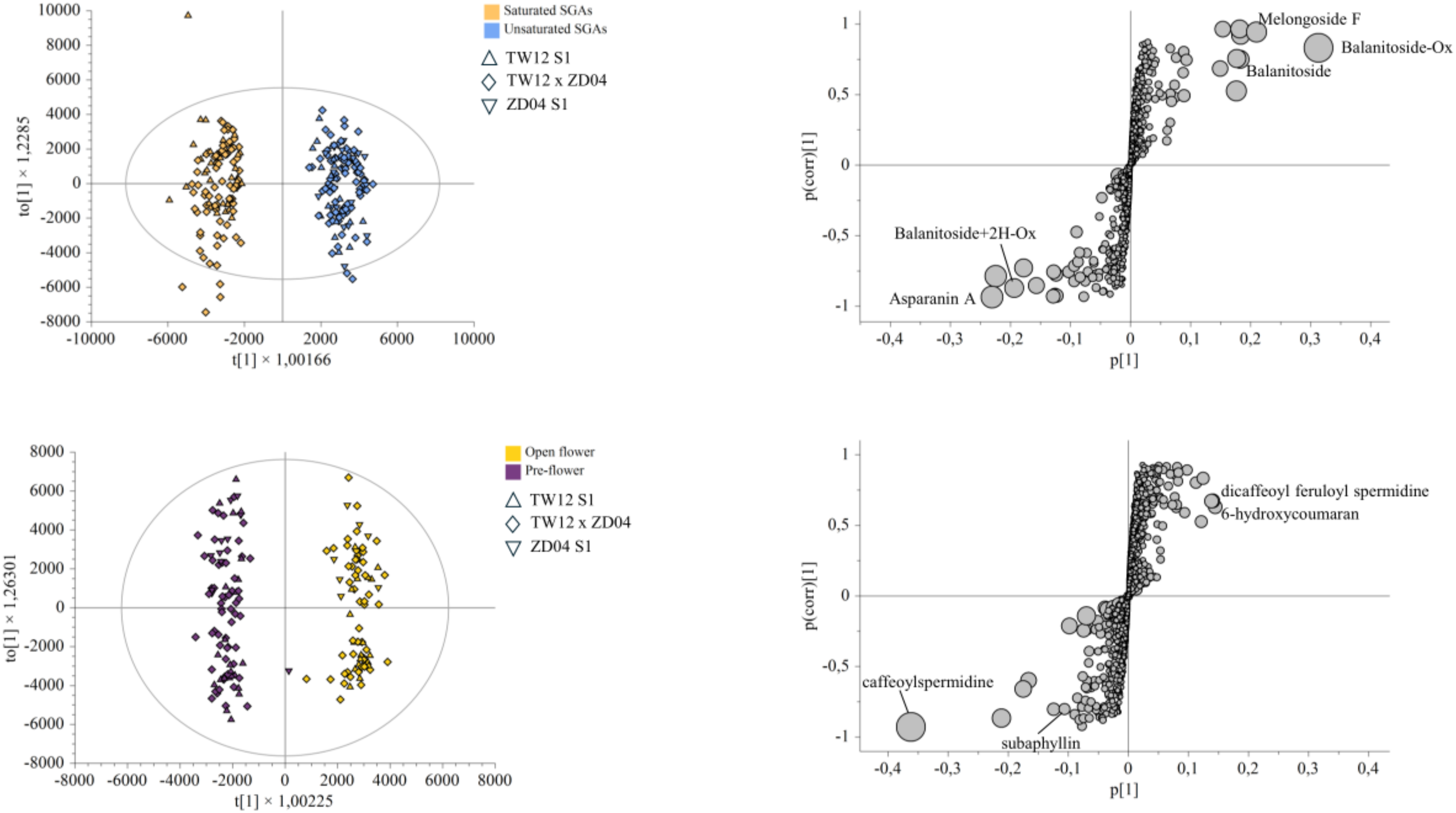

### Fig. S2

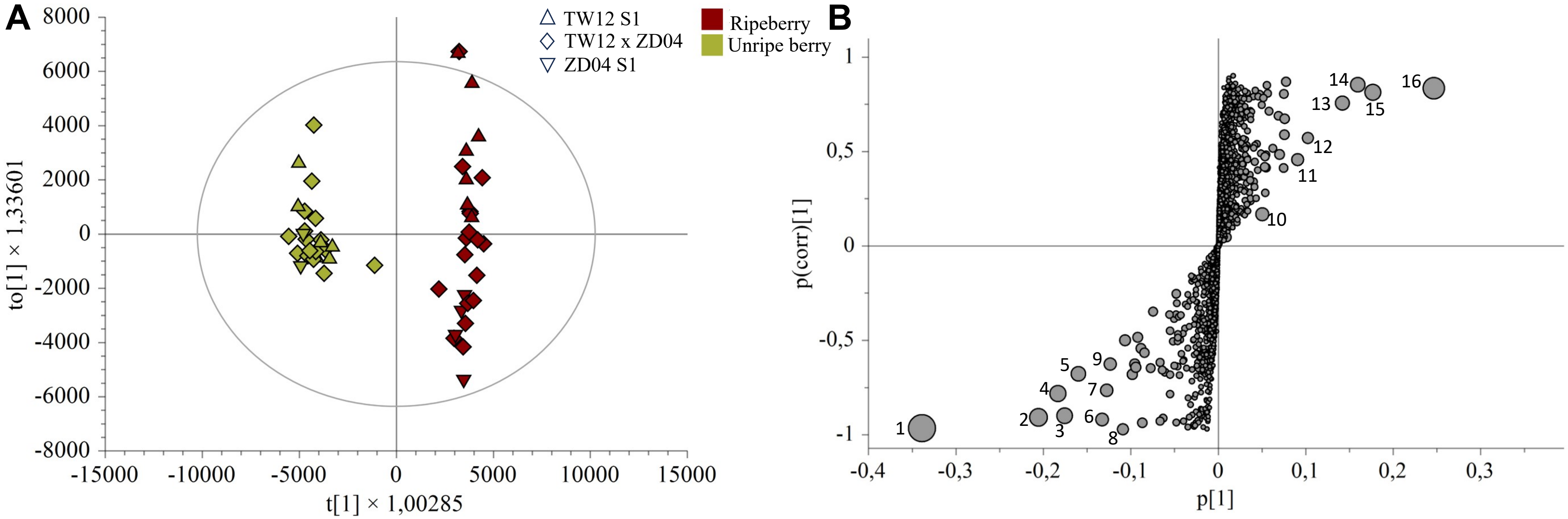
